## Supplementary Table 1 for "Unveiling chemotherapy-induced immune landscape remodeling and metabolic reprogramming in lung adenocarcinoma by scRNA-sequencing"

The table shows the comparative analysis of differences in genetic testing results for all samples in this study.

|  | Control | NCT | p-Value |
| --- | --- | --- | --- |
| EGFR WT | 2 | 1 | 0.524 |
| EGFR Mut | 2 | 4 |  |
| ALK WT | 0 | 3 | 0.167 |
| ALK Tran | 4 | 2 |  |
| PD1 (-) | 3 | 1 | 0.206 |
| PD1 (+) | 1 | 4 |  |
| PDL1 (-) | 2 | 2 | 0.999 |
| PDL1 (+) | 2 | 3 |  |
