## Supplementary Table 2 for "Unveiling chemotherapy-induced immune landscape remodeling and metabolic reprogramming in lung adenocarcinoma by scRNA-sequencing"

Clinical characteristics of patients included in this study.

| Sample | Age | Sex | Stage | Location | Gene mutation | Smoking history | Normal tissue | Tumor tissue | Peripheral blood |
| --- | --- | --- | --- | --- | --- | --- | --- | --- | --- |
| Patient 1 | 60-65 | Female | IB | RUL | EGFR | No | No | Yes | No |
| Patient 2 | 55-60 | Male | IB | LLL | EGFR | No | No | Yes | No |
| Patient 3 | 70-75 | Male | IA | LLL | EGFR | No | No | Yes | No |
| Patient 4 | 65-70 | Female | IIB | RLL | EGFR | No | No | Yes | No |
| Patient 5 | 70-75 | Male | IIIA | LUL | EGFR | Current | No | Yes | No |
| Patient 6 | 75-80 | Male | IA | LUL | EGFR | No | No | Yes | No |
| Patient 7 | 60-65 | Female | IB | RUL | EGFR | Current | No | Yes | No |
| Patient 8 | 55-60 | Female | IA | RML | EGFR | Current | No | Yes | No |
| Patient 9 | 60-65 | Male | IA | LUL | EGFR | No | No | Yes | No |
