## Supplementary figures and images for "Unveiling chemotherapy-induced immune landscape remodeling and metabolic reprogramming in lung adenocarcinoma by scRNA-sequencing"

### Supplementary Figure 1

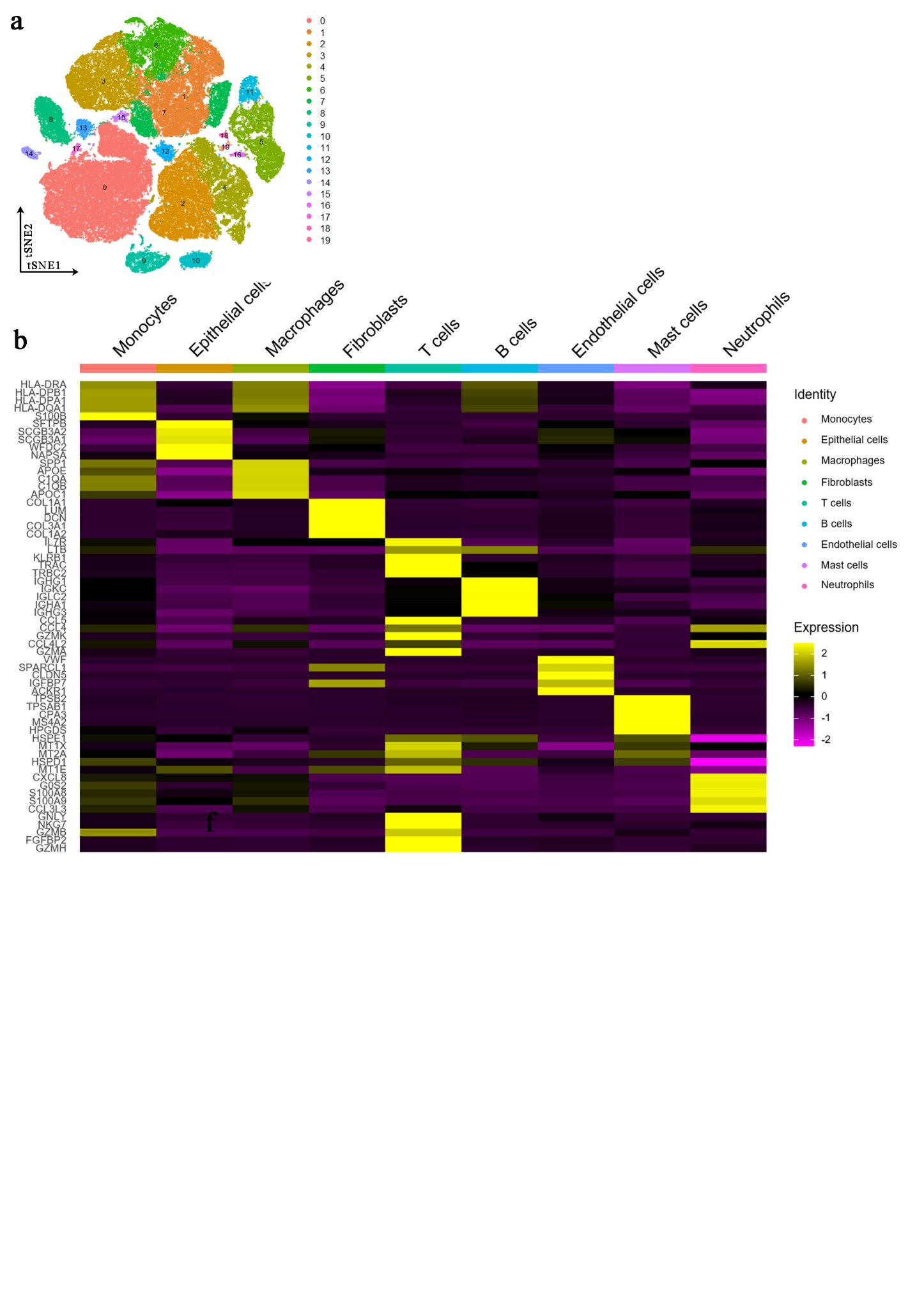

### Supplementary Figure 2

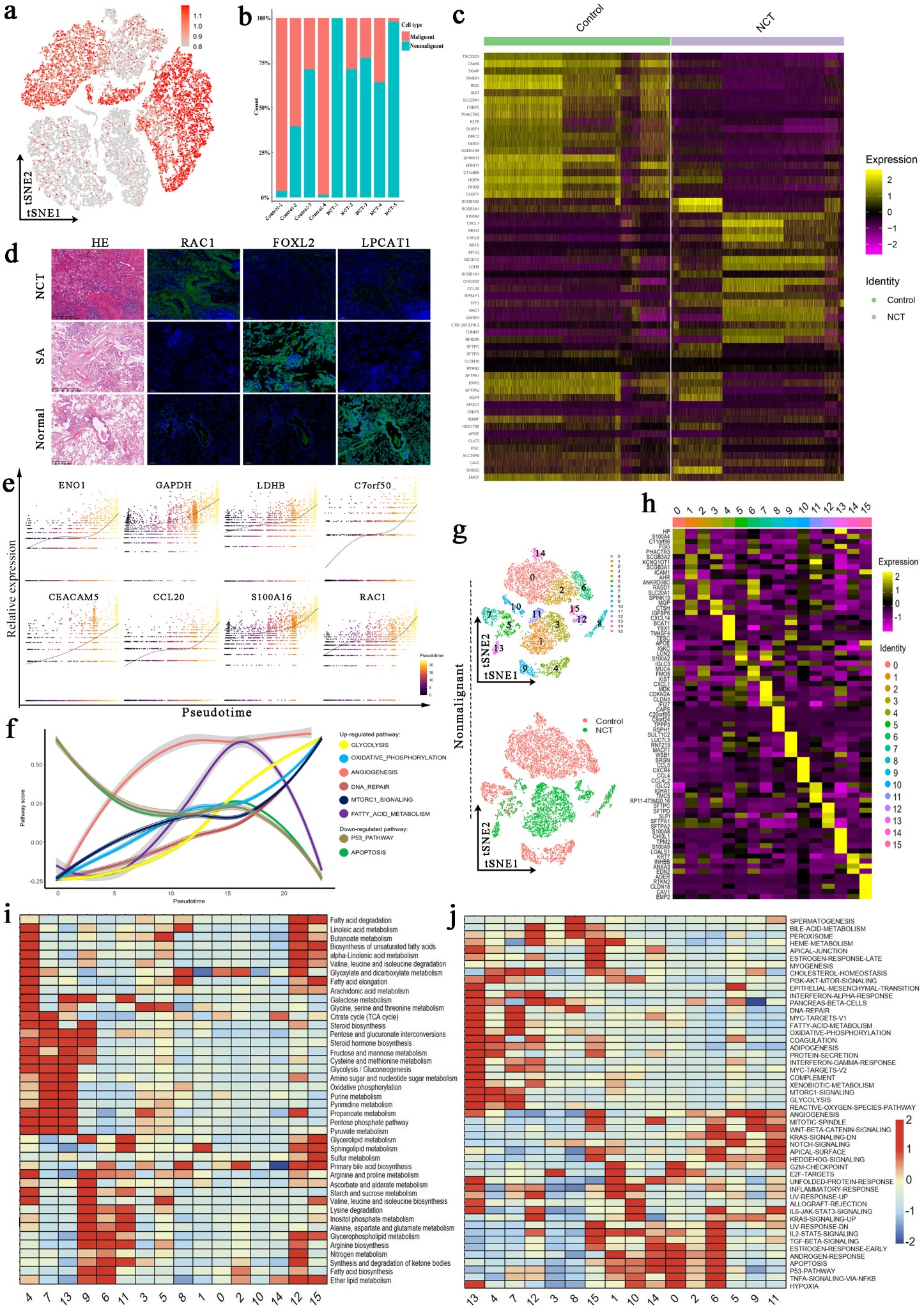

### Supplementary Figure 3

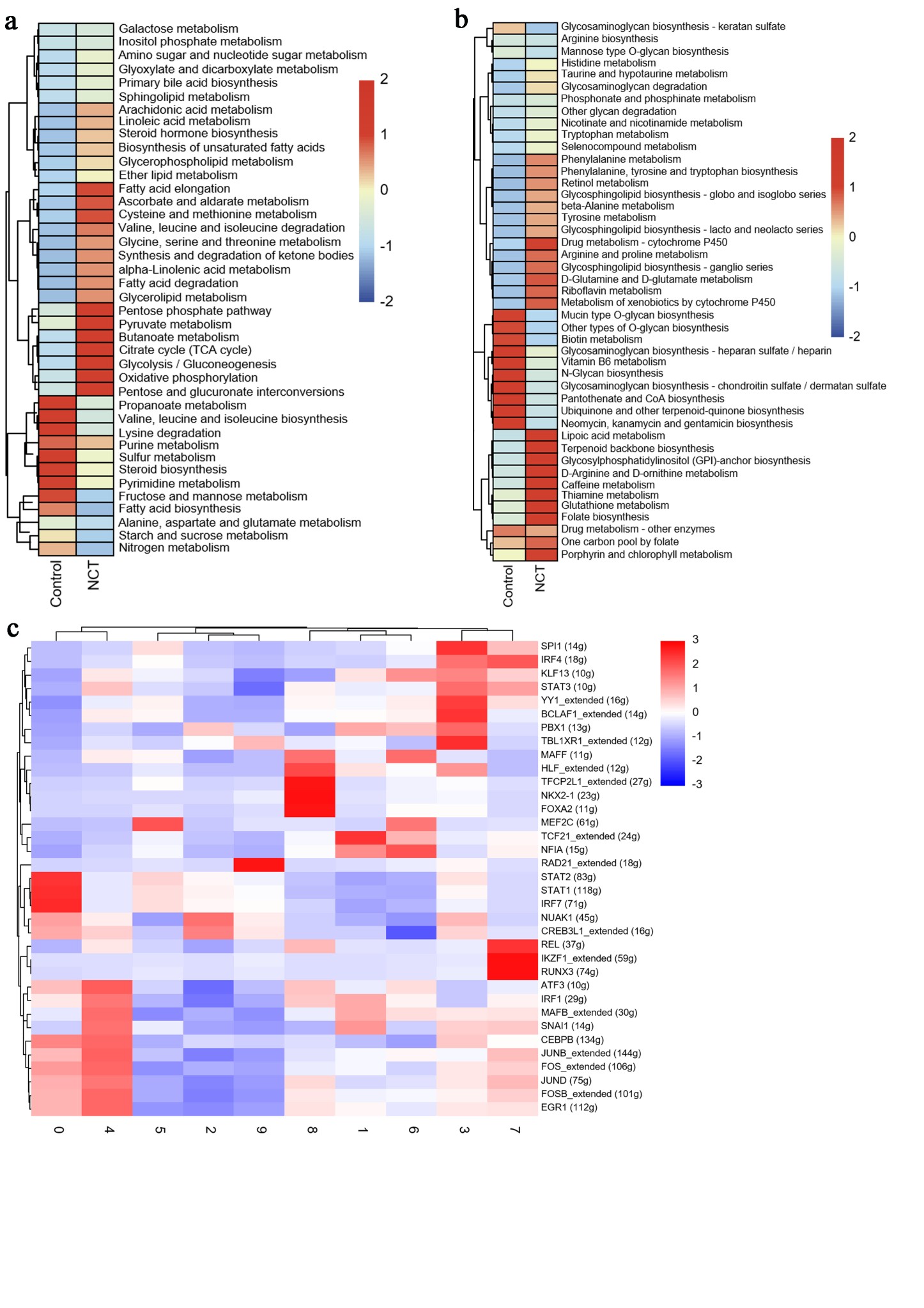

### Supplementary Figure 4

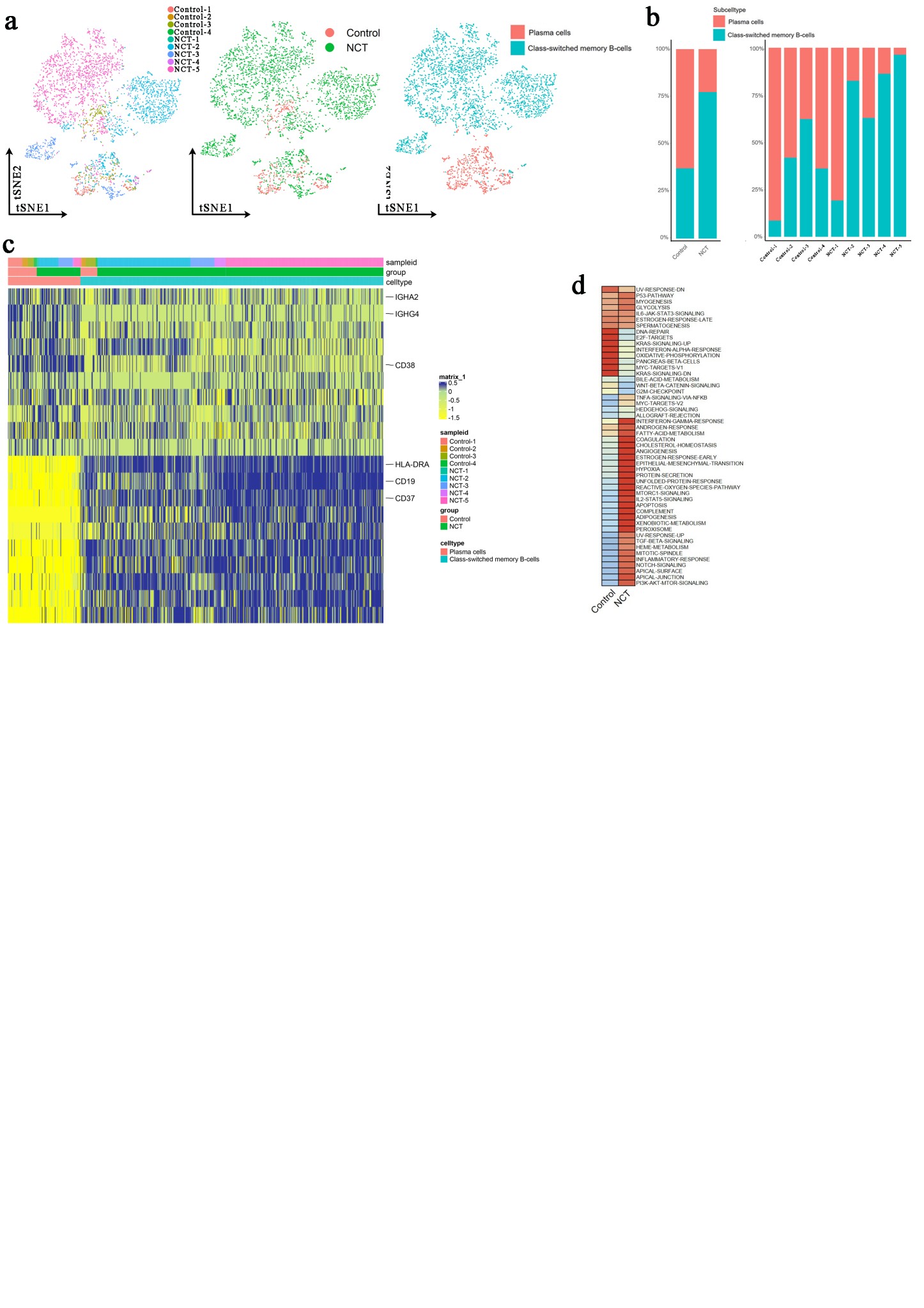

### Supplementary Figure 5

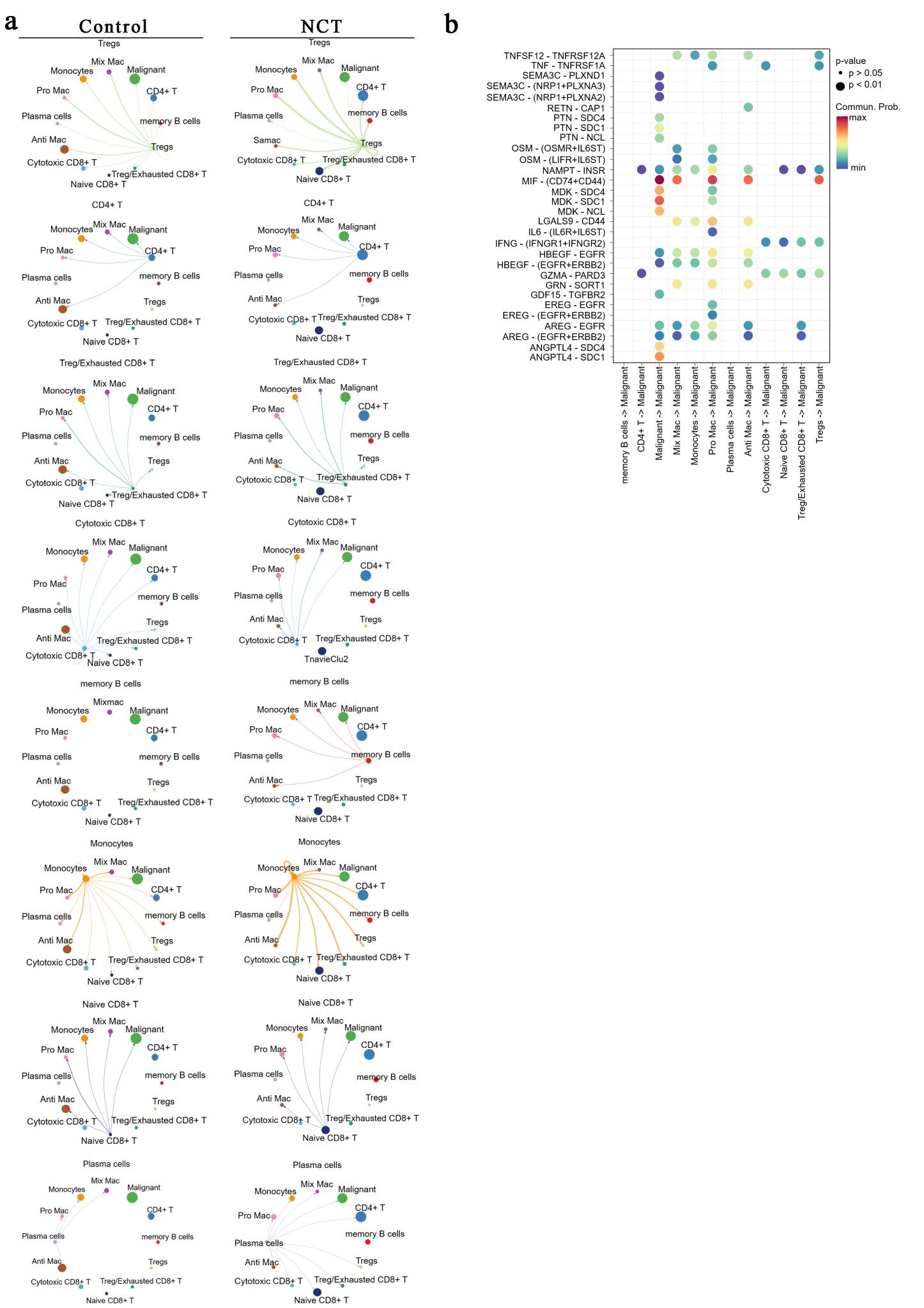

### Supplementary Figure 6

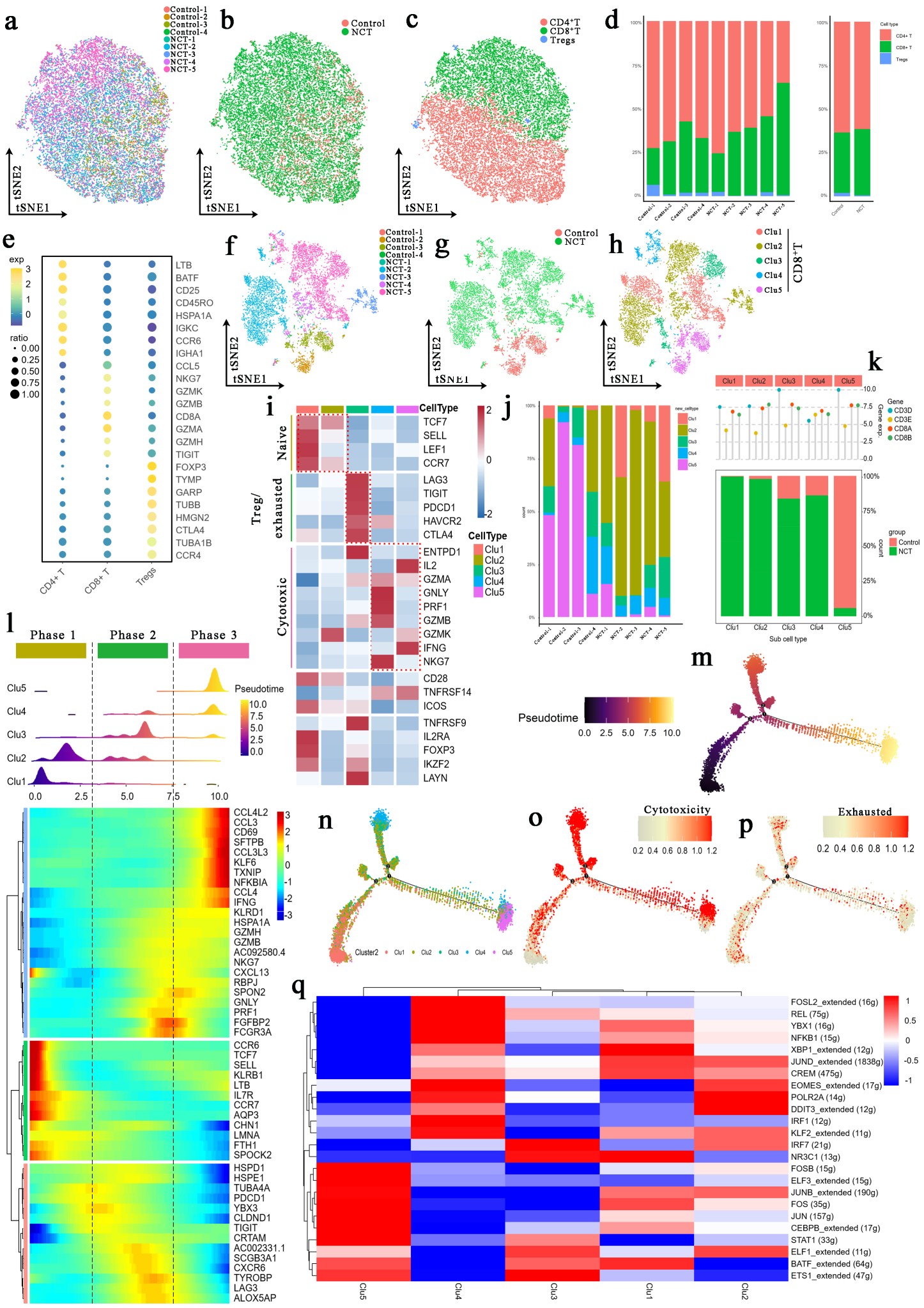

### Supplementary Figure 7

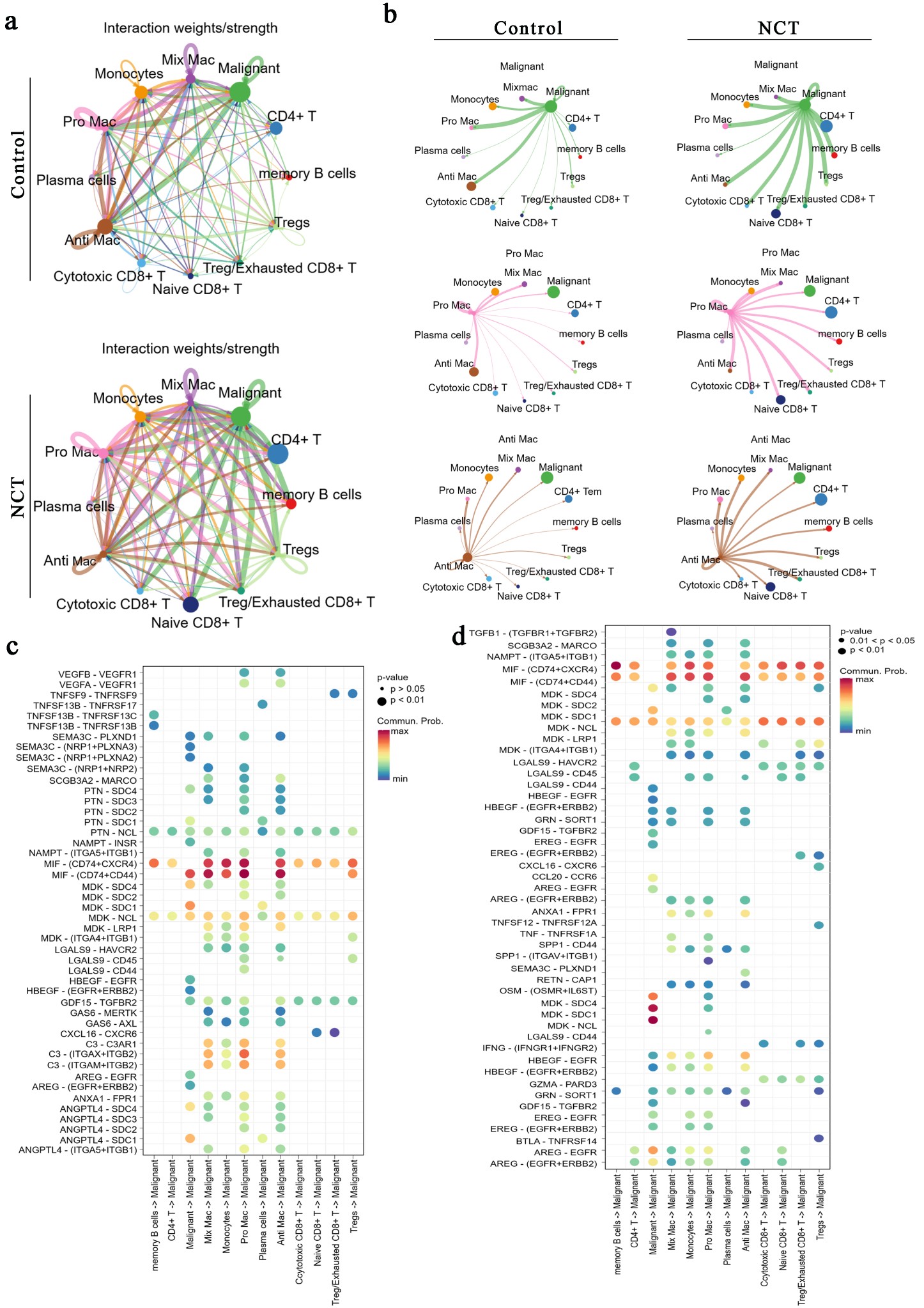
